# Comprehensive analysis of mulberry genetic diversity based on 1-DNJ content and SNP markers

**DOI:** 10.64898/2026.08.11.744330

**Authors:** Zijing Shen, Jiaqi Li, Jianguo Shi, Zekun Li, Fugang Wang, Jin Geng, Kuanlian Hu

**Author notes:** Correspondence authors, (J.-Q. L.); (J.-G. S.).

## Abstract

Mulberry trees have high economic and ecological value, and a robust molecular marker system plus germplasm genetic diversity analysis is critical for innovative utilization of high-quality medicinal and economic mulberry germplasm. Here, 51 mulberry samples were used to develop SNP primers via genome resequencing, with the SNP-PCR system optimized by single-factor and orthogonal assays. The phenotypic diversity and SNP molecular marker genetic diversity of 1-deoxynojirimycin (1-DNJ) in mulberry leaves were analyzed respectively, and the genetic correlation between molecular markers and phenotypic traits was evaluated by Mantel test. Tested germplasm showed marked 1-DNJ variation (0.4805-2.5300 mg/g, CV=0.4241), reflecting rich genetic diversity. The optimal SNP-PCR system included Buffer (containing Mg²^+^) 2.2 μL, 2.5 mM dNTP 0.4 μL, forward and reverse primers (10 μmol·μL^-1^) totaling 2.75 μL, Taq DNA polymerase (5 U·μL^-1^) 0.3 μL, DNA (50 ng·μL^-1^) 1.1 μL, and ddH_2_O 13.65 μL. 23 highly polymorphic ones amplified 91 loci (81 polymorphic, 89.10% polymorphism rate). Genetic diversity analysis showed that the average genetic distance was 0.3010, and the average expected heterozygosity (H) and Shannon information index (I) reached 0.4667 and 0.3104 respectively, indicating that the genetic differentiation among the tested mulberry germplasms was significant and the population had a moderate to upper level of genetic diversity. UPGMA clustering divided 51 germplasms into 6 major groups at a genetic similarity coefficient of about 0.7, while phenotypic clustering based on 1-DNJ content divided them into 2 major categories and 4 subcategories, with high 1-DNJ germplasm clustered independently. Mantel correlation analysis showed that 6 SNP sites were significantly weakly correlated with 1-DNJ content (r < 0.3, p < 0.05), and can be used as candidate molecular markers for subsequent genetic analysis of 1-DNJ content.This study established a stable mulberry SNP-PCR system, Analyze the molecular genetic characteristics of mulberry germplasm and DNJ phenotypic variation rules respectively, and provide basic data for cluster comparison. and provided a scientific basis for marker database improvement, germplasm identification and molecular-assisted breeding.

## Introduction

Mulberry (*Morus alba* L.) is a perennial woody plant belonging to the Moraceae family. It has the advantages of a long history of cultivation, wide adaptability, rich germplasm resources, and the entire plant being medicinally useful. It is currently an important medicinal plant, a raw material for the sericulture industry, and an ecological protection tree species, possessing high economic, ecological, and medicinal comprehensive value[1–3]. Mulberry leaves are praised as the “immortal leaves” in the Compendium of Materia Medica and are rich in flavonoids, polyphenols, alkaloids, and other active ingredients[4]1-Deoxynojirimycin (1-DNJ), a polyhydroxy alkaloid unique to mulberry leaves, is a key substance determining their medicinal value[5, 6]. Research has confirmed that mulberry is currently the species with the highest 1-DNJ content in nature[7–9]. Compared with conventional chemical hypoglycemic drugs, it has the advantages of significant blood sugar-lowering effects and high safety, making it the most promising plant resource for the development of natural hypoglycemic drugs and functional foods[10]. With the global demand for natural medicinal products continuously rising, the efficient evaluation, genetic analysis, and targeted breeding of mulberry germplasm with high 1-DNJ content have become core scientific issues for the sustainable utilization of mulberry genetic resources.

During long-term evolution and the process of artificial cultivation and domestication, mulberry trees have developed rich genetic diversity in agronomic traits and nutrient accumulation due to influences from planting environments, cultivation patterns, and fertilization and irrigation management. At present, traditional molecular markers such as ISSR (Inter-Simple Sequence Repeat), AFLP (Amplified Fragment Length Polymorphism), and SSR (Simple Sequence Repeat) have been widely used to analyze the kinship relationships and genetic diversity of mulberry germplasm, with some studies also conducting association analyses in combination with phenotypic traits and geographic distribution. In previous studies,83 mulberry genotypes were divided into four main clusters using ISSR molecular markers[11]. Similarly, clustering analysis based on morphological traits such as leaf shape, plant height, branch thickness, and internode length also divided them into four main groups, and the study ultimately found that the clustering results based on ISSR molecular markers were generally consistent with the classification results based on morphological traits. DNA barcoding, RAPD, and ISSR techniques have been used to analyze the genetic diversity of black and white mulberries, with SCAR (Sequence Characterized Amplified Region) markers showing a 100% success rate in distinguishing true black and white mulberry samples[12]. SSR molecular markers were used for fingerprinting mulberry varieties from different regions of eastern Anatolia, Turkey, indicating that white mulberry genotypes can serve as valuable resources for mulberry breeding programs[13]. Genetic characteristics of 34 collections of fruit mulberry germplasm were investigated using SRAP (Sequence-Related Amplified Polymorphism) molecular markers, showing clear regional clustering traits[14]. Genetic clustering analysis divided them into five geographic groups. Despite these studies having made progress in analyzing genetic diversity and identifying mulberry germplasm through traditional molecular markers, none have focused on 1-DNJ, a key medicinal trait. Since 1-DNJ content is a typical quantitative trait, it is easily influenced by environmental conditions, growth stages, and cultivation measures, making phenotypic identification alone insufficient to objectively reflect the true genetic differences among germplasm. Moreover, traditional molecular markers such as ISSR, AFLP, and SSR have limitations in terms of low genome coverage and insufficient detection throughput, making them inefficient for matching the needs of genetic analysis of quantitative traits. The combination of these factors has led most existing studies to be limited to either phenotypic analysis of 1-DNJ content alone or studies on molecular genetic diversity. For example, phenotypic analysis focusing only on the 1-DNJ content in mulberry leaves after frost clarifies the optimal harvest time of 1-DNJ in black mulberry and common mulberry, but does not involve genetic-level analysis[15]. ISSR molecular marker technology has been used for systematic genotyping and analysis of Iranian white mulberry (*Morus alba* L.) populations, clarifying the genetic structure characteristics of white mulberry populations in that region[16]; however, during the research process, key medicinal active components such as hydroquinone in white mulberry were not considered, and no association analysis between genotype and medicinal active component content was conducted. Currently, comprehensive genetic analysis combining 1-DNJ content with efficient molecular markers is relatively scarce, with only studies combining 1-DNJ content with high SSR markers [8]. Therefore, there is an urgent need for a molecular marker technology that is widely covering, high-throughput, and highly stable to more comprehensively analyze the genetic basis of 1-DNJ content variation.

Single nucleotide polymorphism (SNP) is a genetic polymorphism formed by the variation of a single nucleotide in a DNA sequence, and it is currently the most widely distributed and densely populated genetic marker[17, 18]. With the advancement of high-throughput technologies, mulberry-specific SNP genotyping chips, Illumina adapter chips, and other genotyping systems can simultaneously detect tens of thousands to millions of SNP markers, offering advantages of high efficiency, low cost, and high automation[19–21]. Currently, SNP technology has been widely used in the mining of SNP loci in mulberry, construction of genetic maps, GWAS (Genome-Wide Association Study) analysis, evaluation of germplasm diversity, and molecular-assisted selection, among other fields[22]. It has become the core technological support for overcoming the aforementioned research difficulties and conducting a comprehensive analysis of 1-DNJ content and molecular genetic diversity, laying a solid foundation for subsequent targeted research.

Based on the shortcomings of the aforementioned research, this study uses 51 mulberry germplasms. Based on the 1-DNJ content measured in leaves, SNP primers were developed from resequencing data, the SNP-PCR reaction system was optimized, and highly polymorphic markers were screened. Diversity analysis and clustering comparison of phenotypes and molecular markers were carried out respectively, and then the correlation between SNP sites and 1-DNJ content was preliminarily evaluated through the Mantel test. The research results are expected to provide phenotypic and molecular basic data for the screening of high-1-DNJ germplasm and the analysis of the genetic mechanism of this trait.

## Materials and methods

### Experimental materials

The experimental materials consisted of 51 mulberry germplasms (Table 1), including 40 cultivated varieties and 11 wild resources, with a wide geographical origin, covering typical materials from major mulberry-producing regions in both the north and south of China, providing a reliable germplasm foundation for subsequent molecular marker development and trait association analysis. All mulberry germplasms were planted in the Mulberry Germplasm Resource Region of the Shaanxi Provincial Academy of Forestry Sciences in Yulin City, Shaanxi Province. Based on the growth performance of the mulberry germplasms, 29 excellent germplasms suitable for cultivation in northern Shaanxi were selected and sent to Beijing Baimai Tech Company for whole-genome resequencing. Provide genomic sequence data for SNP primer development. Four representative germplasms (Hanza No. 3, Jingsang, Heizhenzhu, and Hongguo No. 1) with extremely different 1-DNJ contents were selected for SNP polymorphism primer development and verification.

**Table 1.**
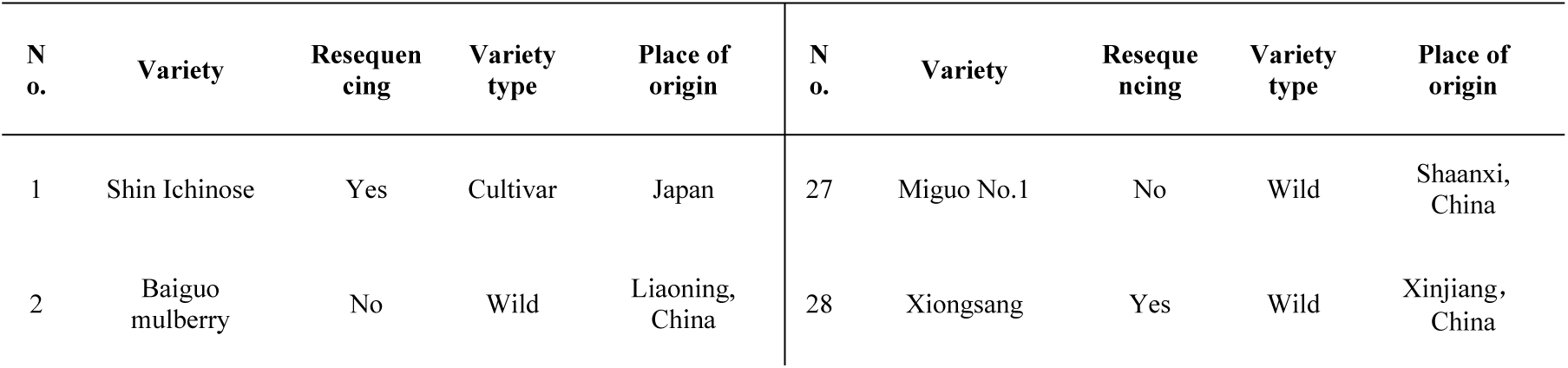
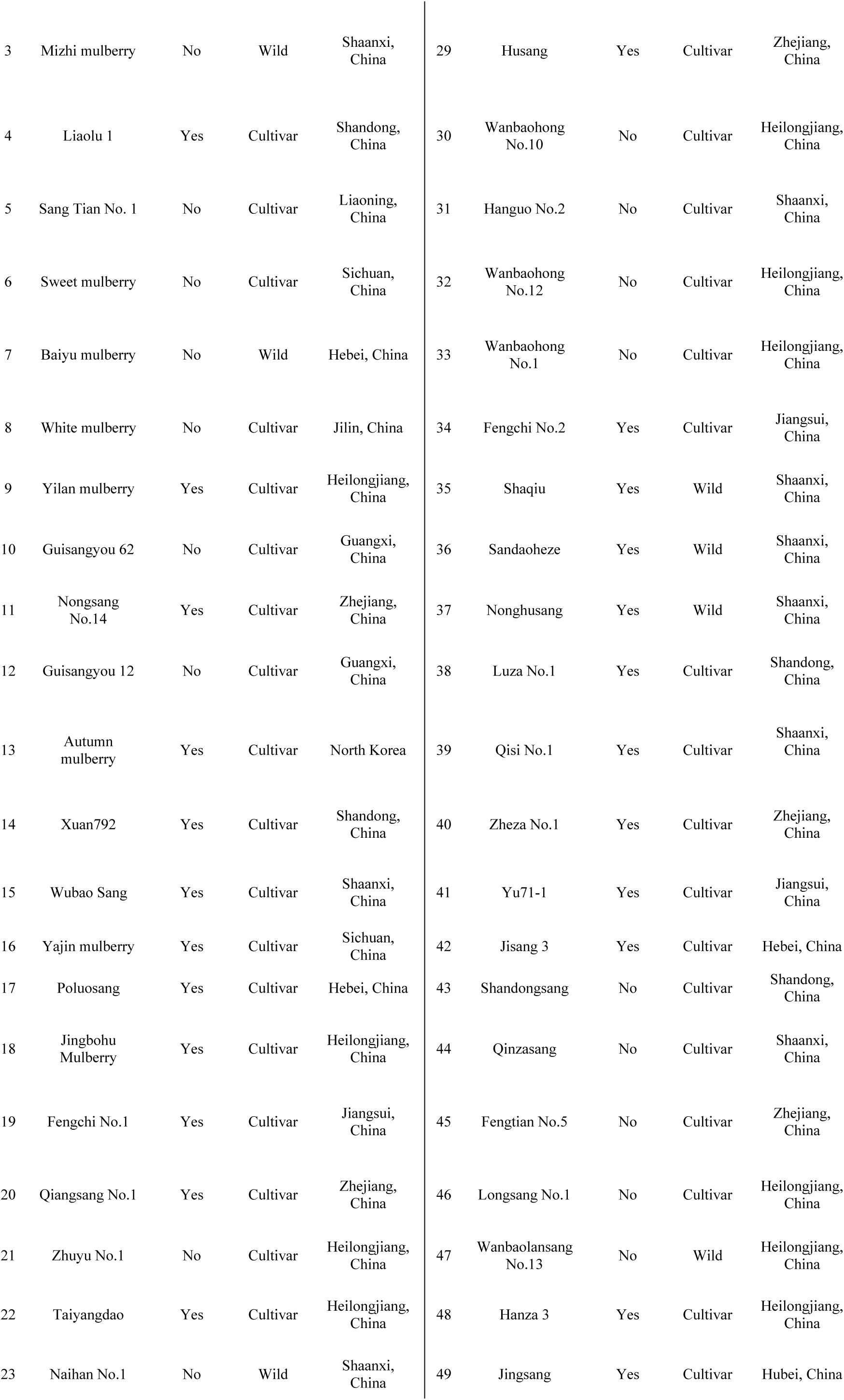
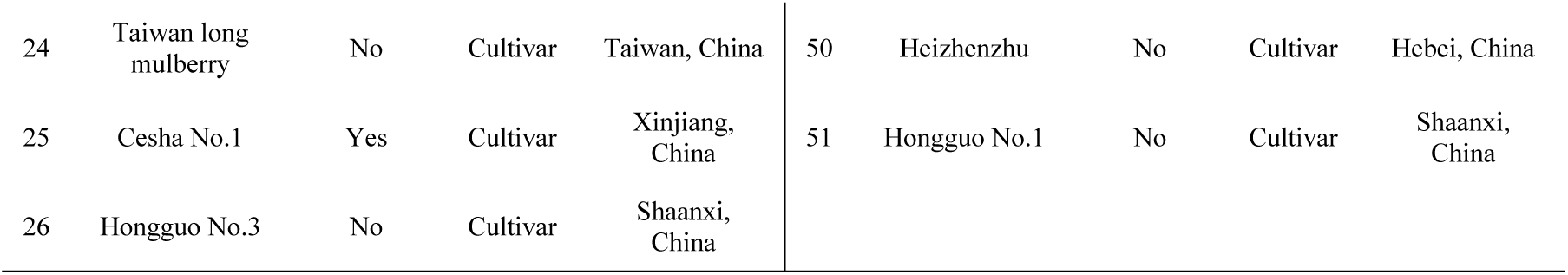
51 mulberry germplasm resources.

| N o. | Variety | Resequencing | Variety type | Place of origin | N o. | Variety | Resequencing | Variety type | Place of origin |
| --- | --- | --- | --- | --- | --- | --- | --- | --- | --- |
| 1 | Shin Ichinose | Yes | Cultivar | Japan | 27 | Miguo No.1 | No | Wild | Shaanxi, China |
| 2 | Baiguo mulberry | No | Wild | Liaoning, China | 28 | Xionsang | Yes | Wild | Xinjiang, China |
| 3 | Mizhi mulberry | No | Wild | Shaanxi, China | 29 | Husang | Yes | Cultivar | Zhejiang, China |
| 4 | Liaolu 1 | Yes | Cultivar | Shandong, China | 30 | Wanbaohong No.10 | No | Cultivar | Heilongjiang, China |
| 5 | Sang Tian No. 1 | No | Cultivar | Liaoning, China | 31 | Hanguo No.2 | No | Cultivar | Shaanxi, China |
| 6 | Sweet mulberry | No | Cultivar | Sichuan, China | 32 | Wanbaohong No.12 | No | Cultivar | Heilongjiang, China |
| 7 | Baiyu mulberry | No | Wild | Hebei, China | 33 | Wanbaohong No.1 | No | Cultivar | Heilongjiang, China |
| 8 | White mulberry | No | Cultivar | Jilin, China | 34 | Fengchi No.2 | Yes | Cultivar | Jiangsui, China |
| 9 | Yilan mulberry | Yes | Cultivar | Heilongjiang, China | 35 | Shaqiu | Yes | Wild | Shaanxi, China |
| 10 | Guisangyou 62 | No | Cultivar | Guangxi, China | 36 | Sandaoheze | Yes | Wild | Shaanxi, China |
| 11 | Nongsang No.14 | Yes | Cultivar | Zhejiang, China | 37 | Nonghusang | Yes | Wild | Shaanxi, China |
| 12 | Guisangyou 12 | No | Cultivar | Guangxi, China | 38 | Luza No.1 | Yes | Cultivar | Shandong, China |
| 13 | Autumn mulberry | Yes | Cultivar | North Korea | 39 | Qisi No.1 | Yes | Cultivar | Shaanxi, China |
| 14 | Xuan792 | Yes | Cultivar | Shandong, China | 40 | Zheza No.1 | Yes | Cultivar | Zhejiang, China |
| 15 | Wubao Sang | Yes | Cultivar | Shaanxi, China | 41 | Yu71-1 | Yes | Cultivar | Jiangsui, China |
| 16 | Yajin mulberry | Yes | Cultivar | Sichuan, China | 42 | Jisang 3 | Yes | Cultivar | Hebei, China |
| 17 | Poluosang | Yes | Cultivar | Hebei, China | 43 | Shandongsang | No | Cultivar | Shandong, China |
| 18 | Jingbohu Mulberry | Yes | Cultivar | Heilongjiang, China | 44 | Qinzasang | No | Cultivar | Shaanxi, China |
| 19 | Fengchi No.1 | Yes | Cultivar | Jiangsui, China | 45 | Fengtian No.5 | No | Cultivar | Zhejiang, China |
| 20 | Qiangsang No.1 | Yes | Cultivar | Zhejiang, China | 46 | Longsang No.1 | No | Cultivar | Heilongjiang, China |
| 21 | Zhuyu No.1 | No | Cultivar | Heilongjiang, China | 47 | Wanbaolansang No.13 | No | Wild | Heilongjiang, China |
| 22 | Taiyangdao | Yes | Cultivar | Heilongjiang, China | 48 | Hanza 3 | Yes | Cultivar | Heilongjiang, China |
| 23 | Naihan No.1 | No | Wild | Shaanxi, China | 49 | Jingsang | Yes | Cultivar | Hubei, China |
| 24 | Taiwan long mulberry | No | Cultivar | Taiwan, China | 50 | Heizhenzhu | No | Cultivar | Hebei, China |
| 25 | Cesha No.1 | Yes | Cultivar | Xinjiang, China | 51 | Hongguo No.1 | No | Cultivar | Shaanxi, China |
| 26 | Hongguo No.3 | No | Cultivar | Shaanxi, China |  |  |  |  |  |

### Mulberry leaf 1-DNJ content detection

On the morning of July 24, 2024, fresh, pest-free mulberry leaves were collected at 9:00–10:00 a.m, placed in preservation bags, the middle leaves were selected as the sampling position. For each accession, three individual plants were chosen, and two leaves were collected from each plant, with a total of six leaves pooled as one sample, and properly labeled. The samples were divided into two portions for processing: 50 g of mulberry leaves were quickly placed in a -80 ℃ freezer for storage, to be used later for leaf genomic DNA extraction and SNP molecular marker analysis. The remaining mulberry leaves were dried in a 60 ℃ constant-temperature oven for 72 hours and then ground into powder to be used for 1-DNJ content determination.

The 1-DNJ content was determined using high-performance liquid chromatography (HPLC). The experiments were carried out using a (Thermo Fisher High Performance Liquid Chromatograph, USA). The specific extraction procedure is as follows: Weigh 0.5 g of the sample, add 25 mL of 0.050 mol/L hydrochloric acid aqueous solution, and assist dissolution with ultrasound for 30 minutes. After cooling to room temperature, centrifuge at 8,000 r/min for 8 minutes and collect the supernatant. The precipitate is extracted once more using the same method, and the supernatants from the two extractions are combined. Adjust the combined supernatant to 50 mL with 0.05 mol/L hydrochloric acid aqueous solution to obtain the crude DNJ extract. The extract was subjected to derivatization, and the 1-DNJ content was determined by high-performance liquid chromatography (HPLC). The average value of the two combined extracts was reported for each sample. The derivatization method and HPLC detection method are carried out according to the reference[16]. The 1-DNJ content in mulberry leaves is calculated using the external standard method. The linear regression equation of the 1-DNJ standard curve is y = 1.655x + 0.264, with a correlation coefficient R²=0.9997, indicating good linearity in the 0.005-0.08 mg/mL concentration range of DNJ, which can be used for quantitative analysis of 1-DNJ content in samples.

### DNA extraction and electrophoresis detection

Take the previously -80°C frozen mulberry leaf samples, cut 0.2 g of young, healthy, pest-free leaves, freeze them in liquid nitrogen, and grind them thoroughly into a fine powder. Extract genomic DNA using the TianGen Biochemical Plant Genomic DNA Extraction Kit (DP305-02, China). Determine DNA purity and concentration via 2% agarose gel electrophoresis(China; 110 V, 25 min) and NanoDropND-2000 (USA) microspectrophotometer, calculating the D260/D280 ratios to select qualified DNA samples. Use a portion of the qualified DNA for library construction and sequencing analysis, dilute the remaining samples to 50 ng·μL^-1^, aliquot, and store at -80°C for subsequent SNP polymorphism primer development and genetic diversity analysis.

### SNP primer design and synthesis

Based on the previous whole-genome resequencing data of 29 mulberry germplasm accessions (PRJNA1304463), SNP sites were screened according to the following criteria: (1) Sites are evenly distributed across the genome; (2) Sites have 100% completeness with no missing information; (3) Sites with a minor allele frequency (MAF) <20% are excluded to ensure basic polymorphism; (4) Sites with a polymorphism information content (PIC) <0.35 are excluded, retaining markers with high information content; (5) Sites that pass the Hardy-Weinberg equilibrium test with a P-value >0.01 are retained to ensure genetic stability; (6) Sites with no other mutations within 50 bp upstream or downstream of the target SNP are selected to avoid interference in subsequent validation experiments. Through the above criteria, high-quality core SNP sites were ultimately obtained, providing reliable targets for subsequent primer design.

Targeting the core SNP sites obtained from screening, SNP primers were designed using Primer 5 software. The primer parameters were set as follows: (1) GC content between 40–60% and annealing temperature between 47 ℃-60 ℃; (2) PCR product of the SNP between 100–400 bp; (3) both forward and reverse primers with lengths of 18– 23 bp. The primers were named according to the rule ‘chromosome number_sequence number’ (e.g. chr1-1F/R). The primers were synthesized (Shenggong Bioengineering Shanghai, China). After synthesis, the primer tubes were centrifuged at 13,000 rpm for 30 s, dissolved to a working concentration of 10 μmol with ddH_2_O according to the instructions, and stored at -80 ℃ for future PCR amplification experiments to provide stable materials.

### Experimental design for SNP-PCR system optimization

Using Baiguo mulberry genomic DNA as a template, primers with strong conservation and stable amplification, Chr2-16 primers, were selected for SNP-PCR system optimization.

### Orthogonal experimental design for SNP reaction system

Determine the optimal system for the SNP reaction, design and conduct an L16(4⁵) orthogonal experiment, selecting five key parameters for optimization: dNTP (2.5 mM), Buffer (containing Mg²+), primers, template DNA, and Taq DNA polymerase (5U). A total of 16 reaction systems were set up, with each combination repeated three times (Table 2).

**Table 2.**
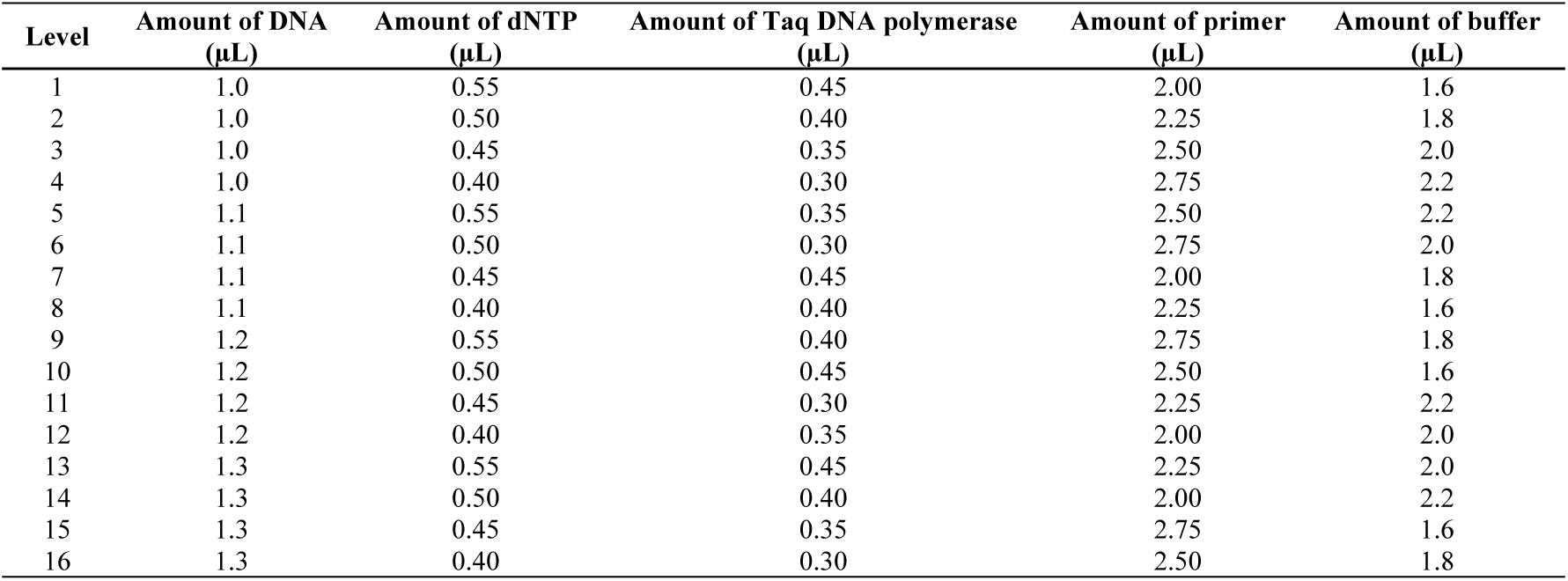
Orthogonal experimental design of SNP reaction.

### SNP reaction system single-factor experimental design

In order to determine the optimal system for the SNP reaction, based on the optimal reaction system determined by the orthogonal experiment, five factors were optimized sequentially: Mg²+ concentration, dNTP concentration, primer concentration, Taq DNA polymerase amount, and template DNA concentration, with two replicates set for each level. During the single-factor optimization process, only the level of the target factor was changed, while the levels of other components in the system remained unchanged (Table 3).

**Table 3.**
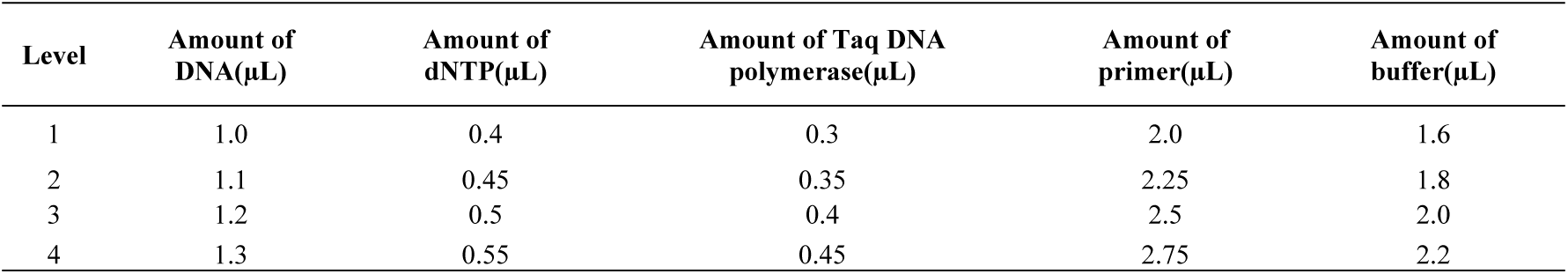
Single-factor experimental design for SNP reaction system.

| Level | Amount of DNA(μL) | Amount of dNTP(μL) | Amount of Taq DNA polymerase(μL) | Amount of primer(μL) | Amount of buffer(μL) |
| --- | --- | --- | --- | --- | --- |
| 1 | 1.0 | 0.4 | 0.3 | 2.0 | 1.6 |
| 2 | 1.1 | 0.45 | 0.35 | 2.25 | 1.8 |
| 3 | 1.2 | 0.5 | 0.4 | 2.5 | 2.0 |
| 4 | 1.3 | 0.55 | 0.45 | 2.75 | 2.2 |

### Validation of the SNP-PCR Optimization System

Based on the optimized reaction system, using the genomic DNA of four mulberry germplasms (Hanza No. 3, Jingsang, Heizhenzhu, and Hongguo No. 1) with extremely different 1-DNJ contents as templates, amplification verification was carried out with the aforementioned optimized PCR reaction system and Chr2-16 primers, the products were analyzed by agarose gel electrophoresis to observe the clarity, specificity, and consistency of the bands.

### SNP-PCR Amplification and Primer Screening

This experiment used the SNP-PCR amplification program as follows: pre-denaturation at 95 ℃ for 3 min; denaturation at 94 ℃ for 30 s, annealing at 58 ℃ for 60 s, extension at 72 ℃ for 60 s, for 30 cycles; final extension at 72 ℃ for 10 min, then stored at 4 ℃. The PCR amplification products were detected using electrophoresis on 2% agarose gel.

Seventy-two pairs of SNP primers (Table 4) were selected, using genomic DNA from Hanza No.3, Jingsang, Heizhenzhu, and Hongguo No.1 as amplification templates. SNP polymorphic primers were screened based on the optimized reaction system. By evaluating the clarity, specificity, stability, and polymorphic performance of the amplified bands, polymorphic primers meeting the requirements were selected. Subsequently, the screened specific primers were used for bulk amplification of DNA from the remaining 47 materials for the genetic diversity analysis of mulberry germplasm.

**Table 4.**
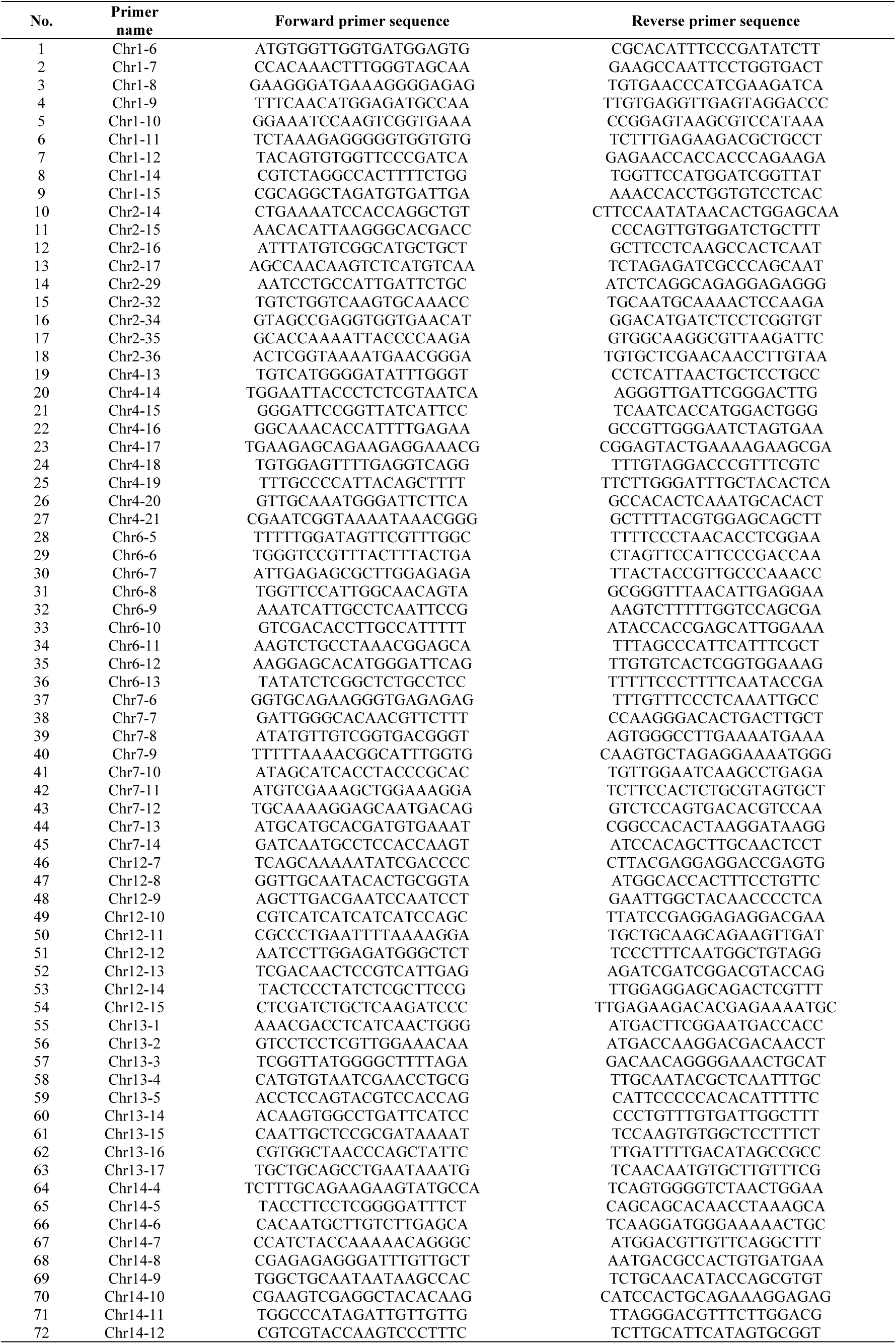
The base sequence of SNP primers.

### SNP Polymorphic Site Statistics and Analysis Processing

The statistical analysis of SNP allele polymorphism sites on the gel was performed using a 0/1 scoring method, where the presence of a band was scored as ‘1’ and the absence as ‘0’; a binary data matrix of 0/1 was constructed using Excel software. PopGen32 software was used to calculate Nei’s diversity (H) of the polymorphic primers. GenAlEx 6.51 software was employed to calculate information such as the observed number of alleles (Na), the effective number of alleles (Ne), Shannon’s information index (I), as well as observed heterozygosity (Ho) and expected heterozygosity (He). NTSYS-pc 2.10e software was used to conduct cluster analysis based on the UPGMA algorithm, construct a cluster dendrogram, and calculate the genetic similarity coefficients and genetic distances (GD) among the test materials. Cluster analysis based on 1-DNJ content was performed using SPSS 27 software. The 51 mulberry germplasm samples were used as indicators, and the detected 1-DNJ content values for these 51 samples were used as data points. The data were centralized, Euclidean distance was used as the clustering metric, and clusters were constructed to create a dendrogram.The Mantel test between SNP loci and 1-DNJ content of mulberry was conducted using the platform (https://cloud.metware.cn/#/user/login).

## Results and analysis

### 1-DNJ content in leaves of 51 mulberry germplasms

The chromatograms of the test samples were compared with the DNJ standard curve (Fig 1) to analyze the 1-DNJ content in the leaves of 51 mulberry germplasms (Table 5). The results showed that the 1-DNJ content in the leaves of the tested mulberry germplasms ranged from 0.4805 mg/g to 2.5300 mg/g, with an average of 1.0049 mg/g and a coefficient of variation as high as 0.4241. This high coefficient of variation directly shows that there is rich genetic diversity in the 1-DNJ content trait in the test population, and the phenotypic differences between individuals are significant, which provides a key phenotypic basis for subsequent genetic analysis and molecular breeding.

**Fig 1.**
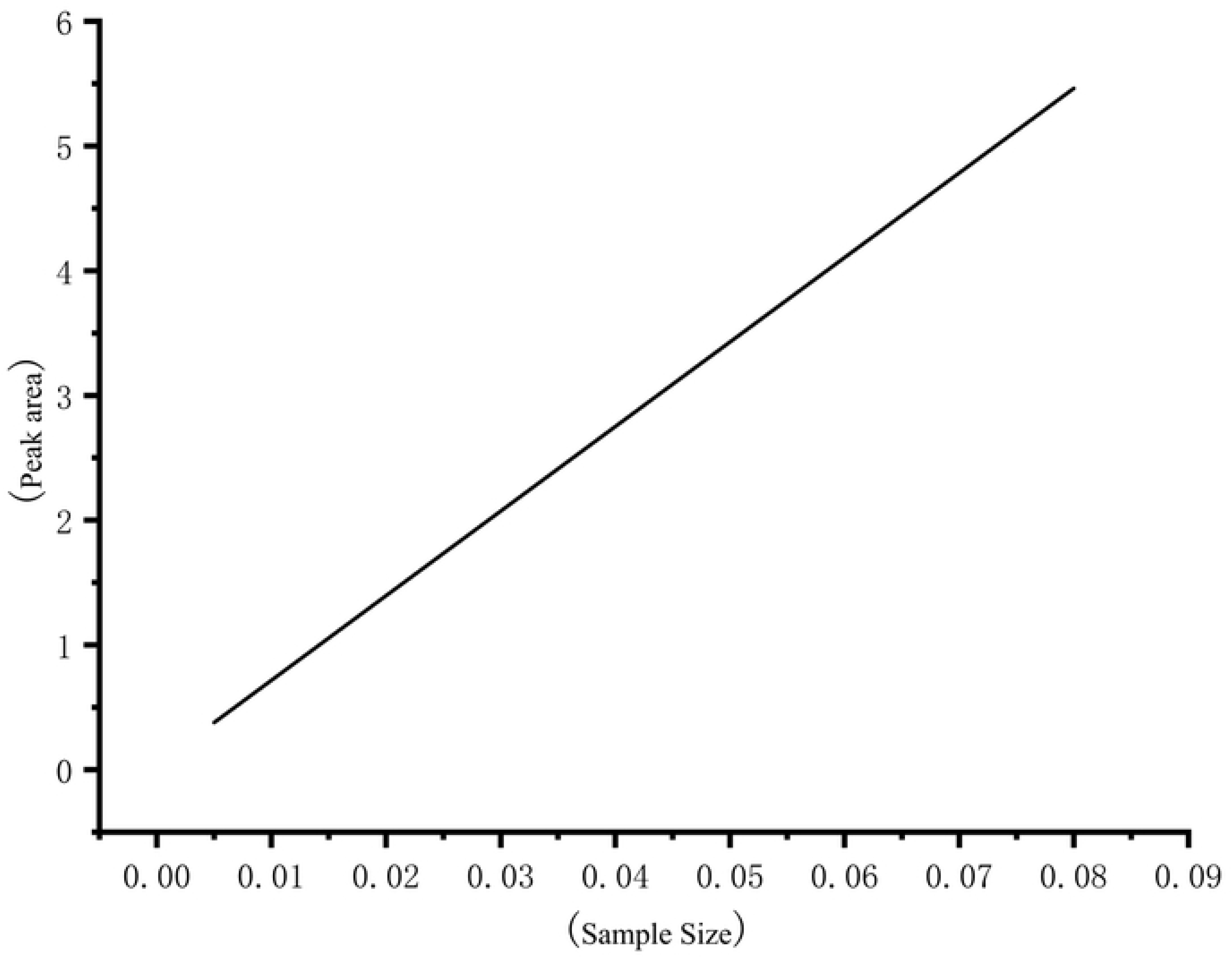
1-DNJ standard curve. **Note:** The linear regression equation was obtained as y = 0.04 + 67.76X, with the correlation coefficient R^2^ = 0.9997.

**Table 5.** 1-DNJ content in mulberry leaves of 51 mulberry germplasm resources.

| No. | Variety | DNJ Content (mg/g) | No. | Variety | DNJContent/ (mg/g) |
| --- | --- | --- | --- | --- | --- |
| 1 | Shin Ichinose | 0.6325 | 27 | Miguo No.1 | 0.8856 |
| 2 | Baiguo mulberry | 0.9666 | 28 | Xionsang | 1.0088 |
| 3 | Mizhi mulberry | 0.9286 | 29 | Husang | 0.9550 |
| 4 | Liaolu 1 | 0.8872 | 30 | Wanbaohong No.10 | 0.8905 |
| 5 | Sang Tian No. 1 | 0.6152 | 31 | Hanguo No.2 | 0.6342 |
| 6 | Sweet mulberry | 0.7930 | 32 | Wanbaohong No.12 | 1.2014 |
| 7 | Baiyu mulberry | 1.0443 | 33 | Wanbaohong No.1 | 0.6357 |
| 8 | White mulberry | 1.1562 | 34 | Fengchi No.2 | 0.9319 |
| 9 | Yilan mulberry | 1.1614 | 35 | Shaqiu | 0.6598 |
| 10 | Guisangyou 62 | 0.9621 | 36 | Sandaohenze | 2.0102 |
| 11 | Nongsang No.14 | 1.3122 | 37 | Nonghusang | 1.1675 |
| 12 | Guisangyou 12 | 0.6317 | 38 | Luza No.1 | 0.5664 |
| 13 | Autumn mulberry | 0.9945 | 39 | Qisi No.1 | 1.0624 |
| 14 | Xuan792 | 0.7657 | 40 | Zheza No.1 | 0.9700 |
| 15 | Wubao Sang | 0.8930 | 41 | Yu71-1 | 2.5300 |
| 16 | Yajin mulberry | 1.5628 | 42 | Jisang 3 | 1.0041 |
| 17 | Poluosang | 0.9038 | 43 | Shandongsang | 0.6557 |
| 18 | Jingbohu Mulberry | 1.9299 | 44 | Qinzasang | 0.9170 |
| 19 | Fengchi No.1 | 0.8657 | 45 | Fengtian No.5 | 1.6012 |
| 20 | Qiangsang No.1 | 1.1973 | 46 | Longsang No.1 | 0.5143 |
| 21 | Zhuyu No.1 | 0.9186 | 47 | Wanbaolansang No.13 | 0.6565 |
| 22 | Taiyangd-ao | 0.8632 | 48 | Hanza 3 | 2.2003 |
| 23 | Naihan No.1 | 1.1254 | 49 | Jingsang | 1.2701 |
| 24 | Taiwan long mulberry | 0.5598 | 50 | Heizhenzhu | 0.6392 |
| 25 | Cesha No.1 | 0.7227 | 51 | Hongguo No.1 | 0.8070 |
| 26 | Hongguo No.3 | 0.4854 |  |  |  |

Among the tested mulberry germplasm resources, the germplasm with the highest 1-DNJ content was Yu 71-1 (2.5300 mg/g), while the one with the lowest content was Hongguo No. 3 (0.4854 mg/g). In terms of content distribution, 16 mulberry germplasms had 1-DNJ content below 0.8 mg/g, including Shin Ichinose, Sang Tian No.1, Sweet mulberry and others; 29 accessions had a 1-DNJ content ranging from 0.8 to 1.5 mg/g, including Baiguo mulberry, Mizhi mulberry, Liaolu No.1 and others; six accessions had a 1-DNJ content above 1.5 mg/g, among which three germplasms (Sandaoheze, Yu71-1, Hanza No.3) showed a 1-DNJ content exceeding 2.0 mg/g. This significant phenotypic differentiation not only confirms the rich genetic diversity of the population but also provides a phenotypic basis for subsequent correlation analysis of 1-DNJ content with SNP markers.

### DNA purity and concentration testing

The gel electrophoresis results of genomic DNA from 51 mulberry germplasm samples showed that the electrophoresis bands were generally clear and bright, with no protein or RNA contamination and no trailing, indicating that the DNA was undegraded and had good integrity (Fig 2). The results from the ultra-micro spectrophotometer (NanoDropND-2000) showed that DNA concentrations ranged from 57.6 to 112.7 ng·μL^-^¹, with an average of 82.3 ng·μL^-^¹. The D260/D280 ratio ranged from 1.58 to 1.98, indicating that both the quality and concentration of DNA met the requirements of this experiment.

**Fig 2.**
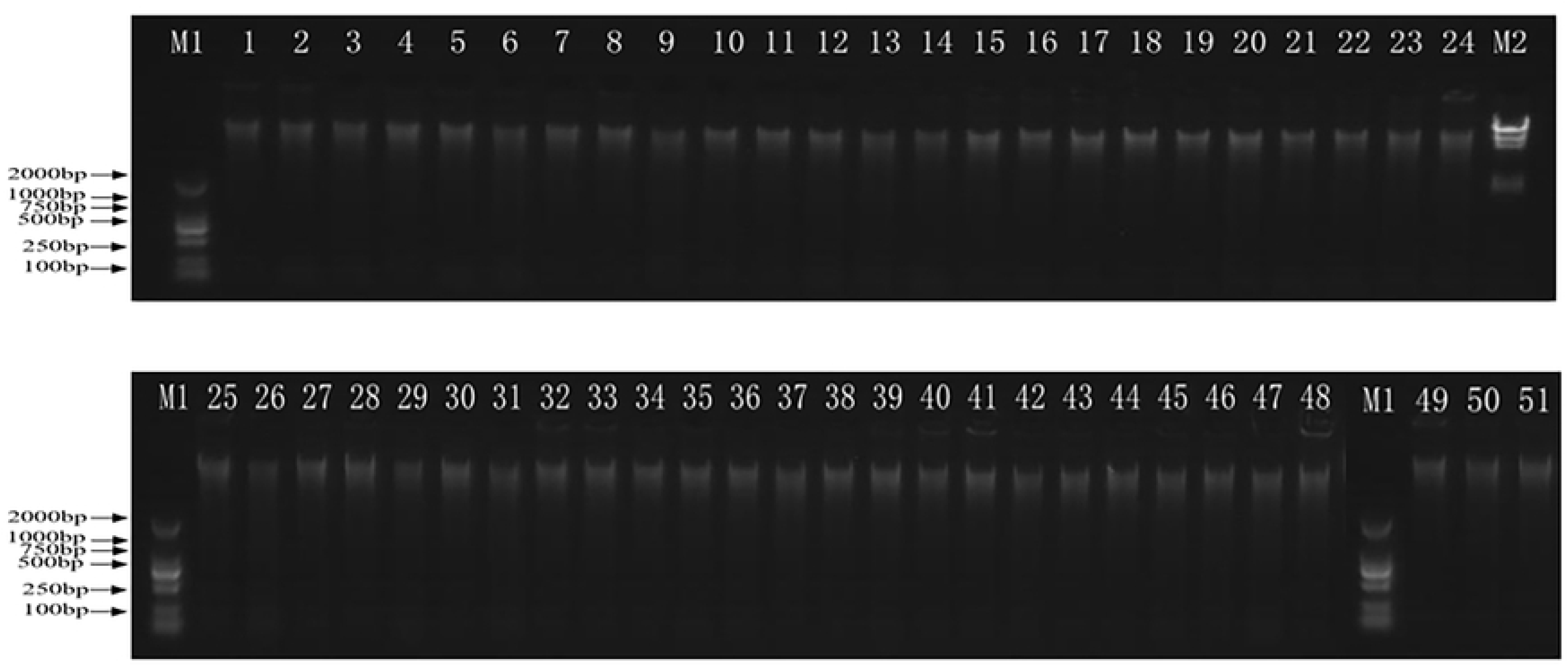
Electrophoretogram of genomic DNA from 51 mulberry germplasms. **Note:** M: 2000 bp DNA Marker; DNA activity detection of 51 mulberry trees, with DNA sequences identical to those in (Table 1)

### Orthogonal test tesults of the mulberry SNP-PCR reaction system

Screen the SNP-PCR reaction system based on L₁₆(4⁵) orthogonal design. The amplification effects of the 16 combinations were significantly different (Fig 3), and most combinations had problems such as no amplification, non-specific bands, or poor repeatability. Among them, the amplification band of the fourth combination is clear and complete, and the results are stable and consistent three times, and the effect is the best. Therefore, the fourth combination was selected as the optimal system for the orthogonal experiment. Its 20 μL reaction composition is: 2.2 μL of Buffer (containing Mg²⁺), 0.4 μL of 2.5 mM dNTP, 2.75 μL of upstream and downstream primers (10 μmol·μL⁻¹), 0.3 μL of Taq DNA polymerase (5 U·μL⁻¹), and template DNA (50 ng·μL⁻¹) 1.0 μL, ddH₂O 13.35 μL.

**Fig 3.**
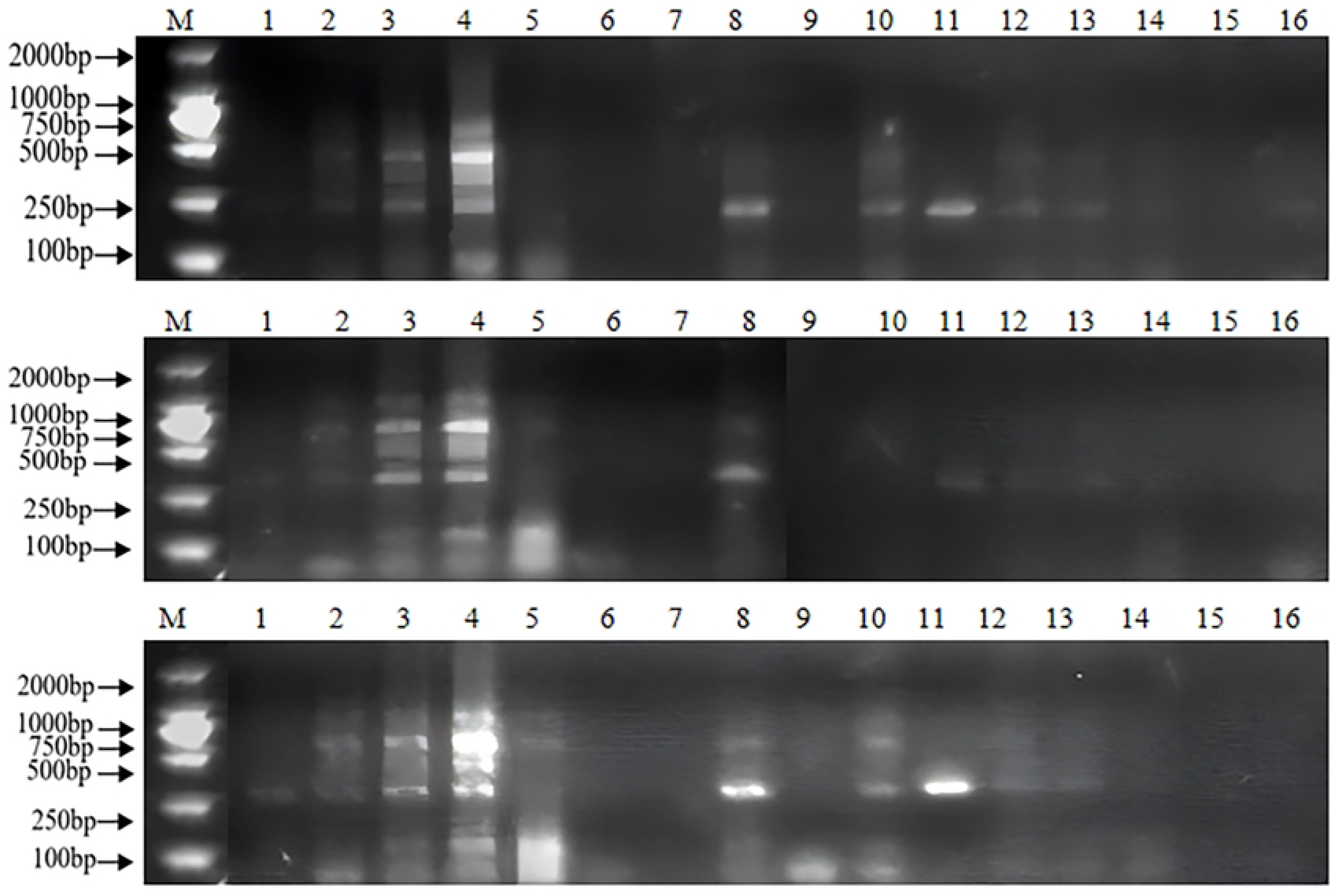
Amplification results of 16 treatments in mulberry SNP orthogonal experiment. **Note:** M: 2000 bp DNA Marker; Shows the results of three replicates of the orthogonal experiment.

### Single-factor test results of the mulberry SNP reaction system

Based on the optimal combination 4 screened via orthogonal experiments, the dosage of each component was further optimized through single-factor experiments. Based on the brightness, clarity and repeatability of the amplified band (Fig 4), the optimal dosage of each component was determined to be: template DNA 1.1 μL, dNTP (2.5 mM) 0.4 μL, Taq DNA polymerase (5 U·μL⁻¹) 0.3 μL, up/downstream primer (10 μmol·μL⁻¹) 2.75 μL, Buffer (containing Mg²⁺) 2.2 μL. Except for the fine adjustment of the template DNA dosage from 1.0 μL to 1.1 μL, the optimization levels of the other factors were consistent with the results of the orthogonal experiment.

**Fig 4.**
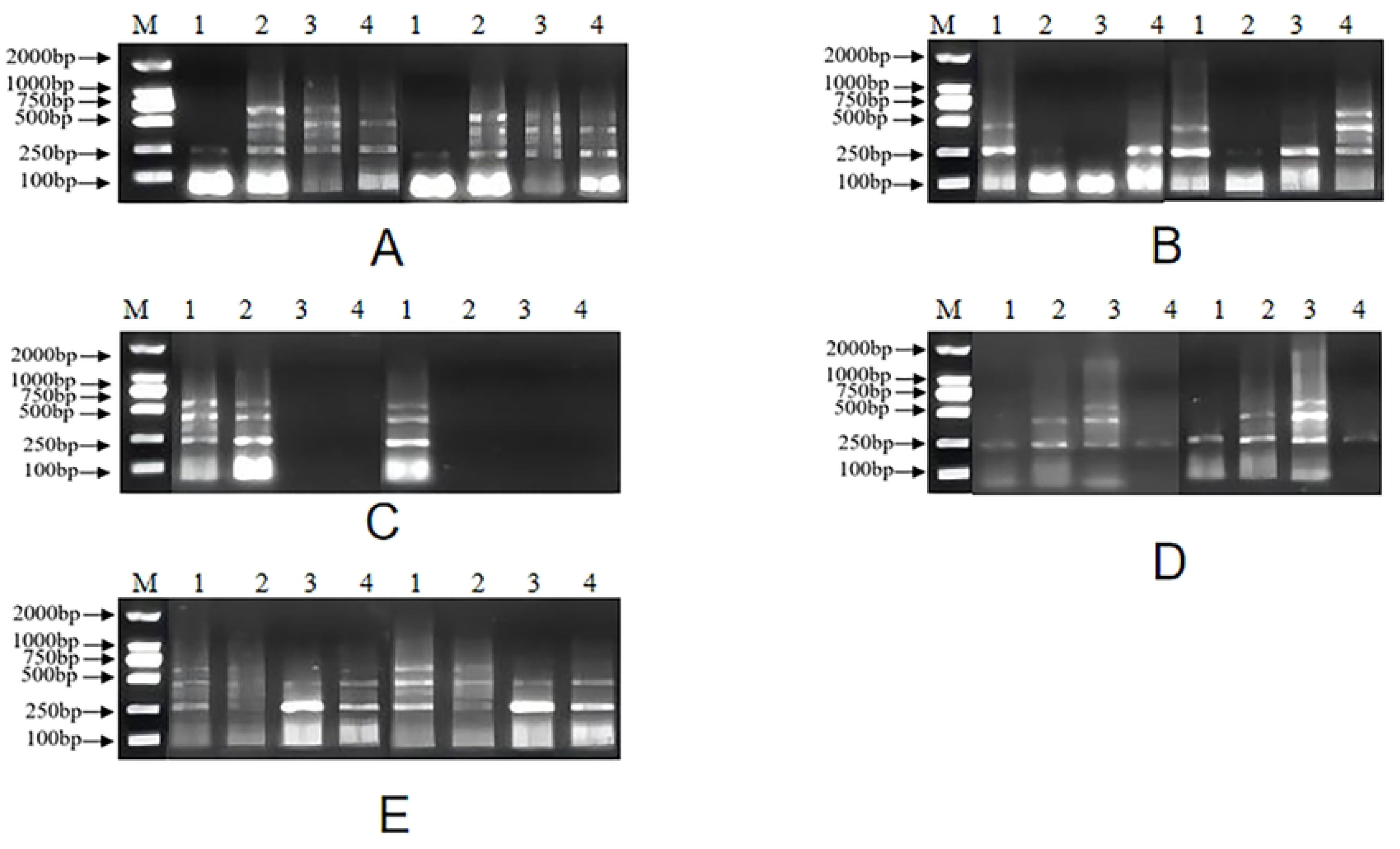
Gel electrophoresis results of SNP-PCR products under different concentration treatments of single factors in mulberry. **Note:** M: 2000 bp DNA Marker; (A) Genomic DNA dosage (Lanes 1–8 correspond to two replicates of Levels 1–4 in Table 4, respectively: 1.0 μL, 1.1 μL, 1.2 μL, 1.3 μL); (B) dNTP dosage (Lanes 1–8 correspond to Levels 1–4 in Table 4, respectively: 0.4 μL, 0.45 μL, 0.5 μL, 0.55 μL); (C) Taq DNA polymerase dosage (Lanes 1–8 correspond to Levels 1–4 in Table 4, respectively: 0.3 μL, 0.35 μL, 0.4 μL, 0.45 μL); (D) Primer dosage (Lanes 1–8 correspond to Levels 1–4 in Table 4, respectively: 2.0 μL, 2.25 μL, 2.5 μL, 2.75 μL); (E) Mg²⁺-containing Buffer dosage (Lanes 1–8 correspond to Levels 1–4 in Table 4, respectively: 1.6 μL, 1.8 μL, 2.0 μL, 2.2 μL).

The results show that the orthogonal design can determine the interaction of each factor, and the single-factor experiment can intuitively analyze the impact of each factor level. Except for the amount of template DNA, the optimal levels of the other factors are consistent with the orthogonal results. Comprehensive determination of the optimal reaction system for mulberry SNP-PCR: Buffer 2.2 μL, 2.5 mM dNTP 0.4 μL, upstream and downstream primers (10 μmol·μL⁻¹) 2.75 μL each, Taq DNA polymerase (5 U·μL⁻¹) 0.3 μL, template DNA (50 ng·μL⁻¹) 1.1 μL, add ddH₂O to 20 μL.

### Validation of the optimized SNP-PCR reaction system for mulberry

Based on the previously optimized optimal mulberry SNP-PCR amplification system, the stability and applicability of the system were verified using genomic DNA from four mulberry germplasm accessions. The results demonstrated that all 72 pairs of SNP primers could be stably amplified under this system. Gel electrophoresis analysis showed stable, clear bands with abundant polymorphism (Fig 5), confirming that this amplification system possesses high repeatability and reliability, and can be widely applied to SNP marker analysis of mulberry genomic DNA.

**Fig 5.**
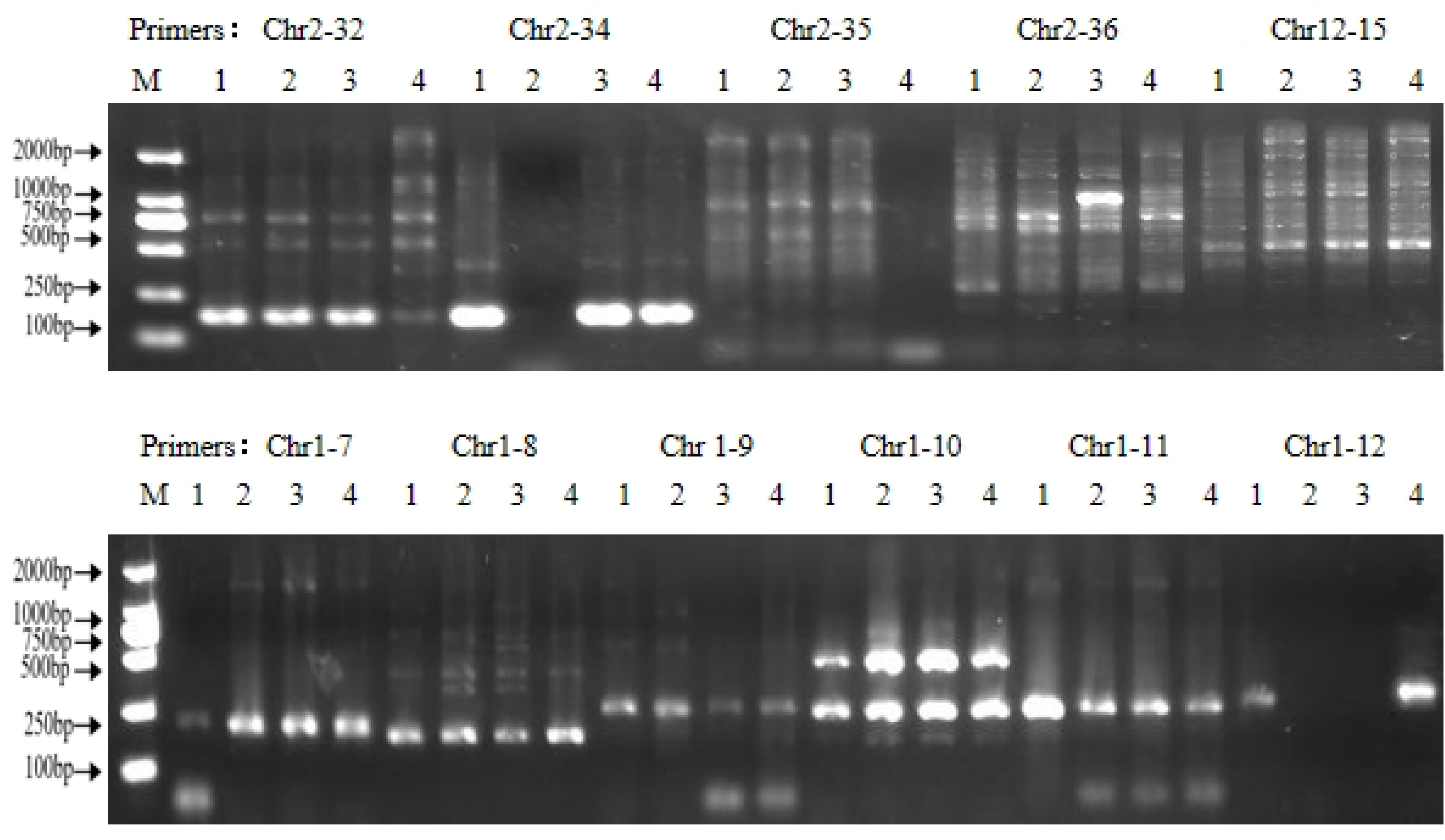
Verification of the optimal SNP-PCR system by partial primer pairs. **Note:** M: 2000 bp DNA Marker; Figure 5 shows selected polymorphic primers used in the experiment, listed in order as Chr2-32, Chr2-34, Chr2-35, Chr2-36, Chr12-15, Chr1-7, Chr1-8, Chr1-9, Chr1-10, Chr1-11, and Chr1-12.

### Genetic diversity analysis of mulberry germplasm resources based on SNP markers

Based on the optimized SNP-PCR system, PCR amplification was performed on 72 pairs of primers using genomic DNA from four mulberry materials as templates. From these, 23 primer pairs with high polymorphism, clear bands and good reproducibility were selected for mulberry genetic diversity analysis. A total of 91 loci were amplified by the 23 primer pairs, of which 81 were polymorphic loci, with an average of 3.52 polymorphic loci per primer pair and a polymorphic locus percentage of 89.10%, indicating abundant genetic diversity in mulberry. For the 23 primer pairs, the observed number of alleles (Na) was 2 for all pairs; the effective number of alleles (Ne) ranged from 1.2378 to 1.9899, with an average of 1.5362; Nei’s genetic diversity index (H) ranged from 0.2947 to 0.3906, with an average of 0.4667; and Shannon’s information index (I) ranged from 0.1881 to 0.4975, with an average of 0.3104. It shows that the SNP primers developed in this study have good polymorphism and can be effectively used for genetic diversity analysis of mulberry germplasm resources (Table 6).

**Table 6.**
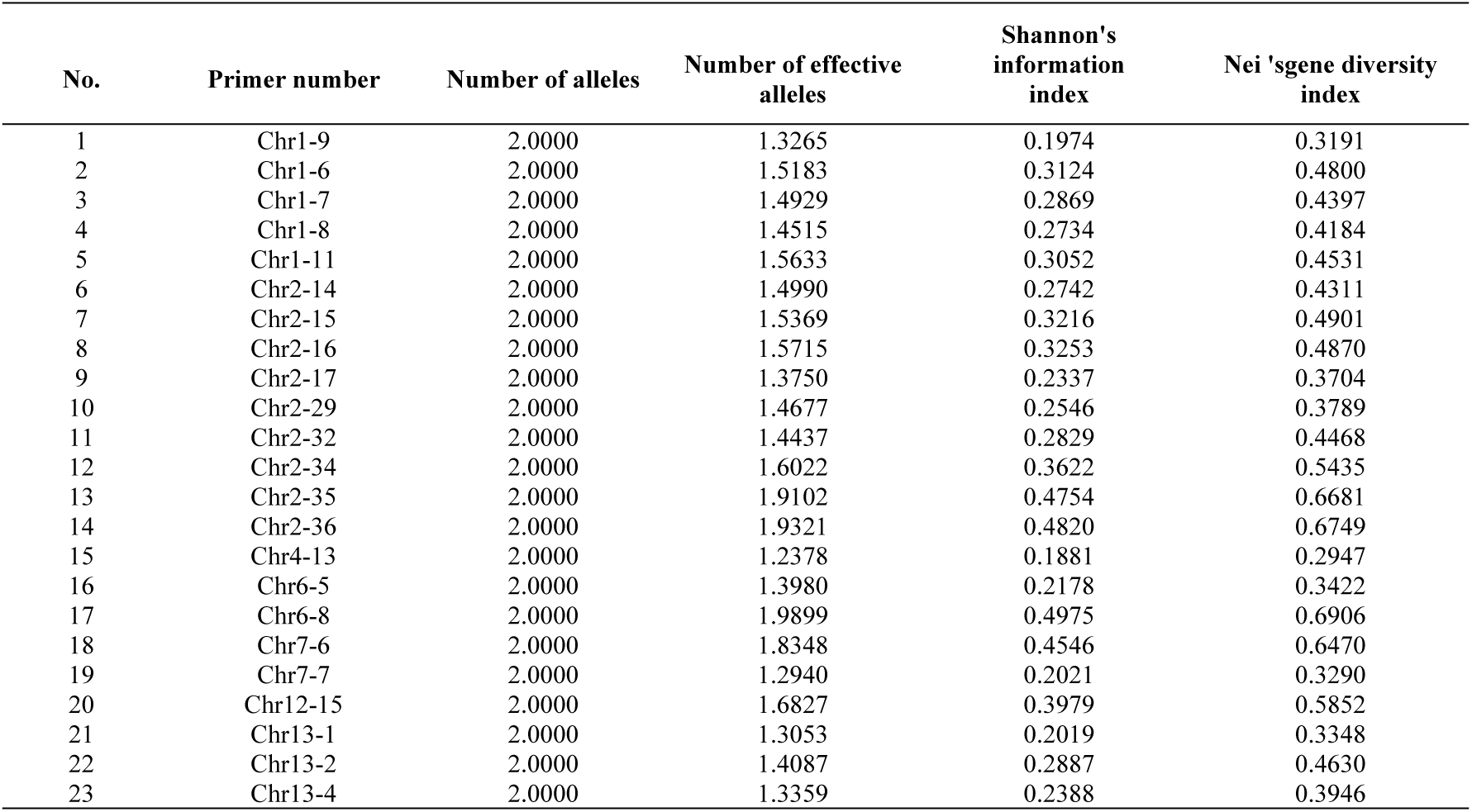
Genetic diversity analysis of 23 primers.

### Genetic distance and cluster analysis of mulberry germplasm resources

Based on the 0/1 matrix of polymorphic sites amplified by 23 pairs of SNP primers, the genetic distance (GD) between 51 mulberry germplasm resources was calculated. The results showed that the genetic distance ranged from 0.2340 to 0.7273, and the average genetic distance was 0.4584, indicating that there was a moderate degree of genetic difference between the materials, and that the core of genetic differentiation was driven by differences in the germplasm’s own genetic background and breeding direction. Among them, Fengchi No.2 and Hongguo No.1 have the smallest GD value (0.2340). The genetic similarity between the two is high, which is due to the significant homology of their genetic backgrounds. They are cultivated germplasm under the same breeding system, and the breeding direction is consistent, and no significant genomic locus differentiation has occurred. Xiongsang and Husang have the largest GD value (0.7273), and there is significant genetic differentiation between germplasms. It is speculated that this may be related to the source of germplasm, ecotype differences and breeding history. Husang is a cultivated germplasm that has been artificially selected for a long time. Xiongsang mostly originates from wild or semi-wild states. The two are driven by differential artificial selection pressure during the process of domestication and adaptation, resulting in significant genetic differentiation at the whole genome level. Based on the 0/1 matrix constructed from 81 polymorphic bands amplified by 23 pairs of primers, UPGMA cluster analysis was performed through NTSYS software (Fig 6). The results showed that the genetic correlation coefficient of 51 mulberry germplasms ranged from 0.61 to 0.89, and the correlation coefficient of the cluster analysis was r = 0.78606, indicating that the clustering results were reliable. At the genetic coefficient of nearly 0.7, 51 mulberry trees can be divided into 6 groups. The first category contains 36 accessions, the second category contains 5 accessions, the third category contains 3 accessions, the fourth category contains 3 accessions, the fifth category contains 2 accessions, and the sixth category contains 2 accessions.

**Fig 6.**
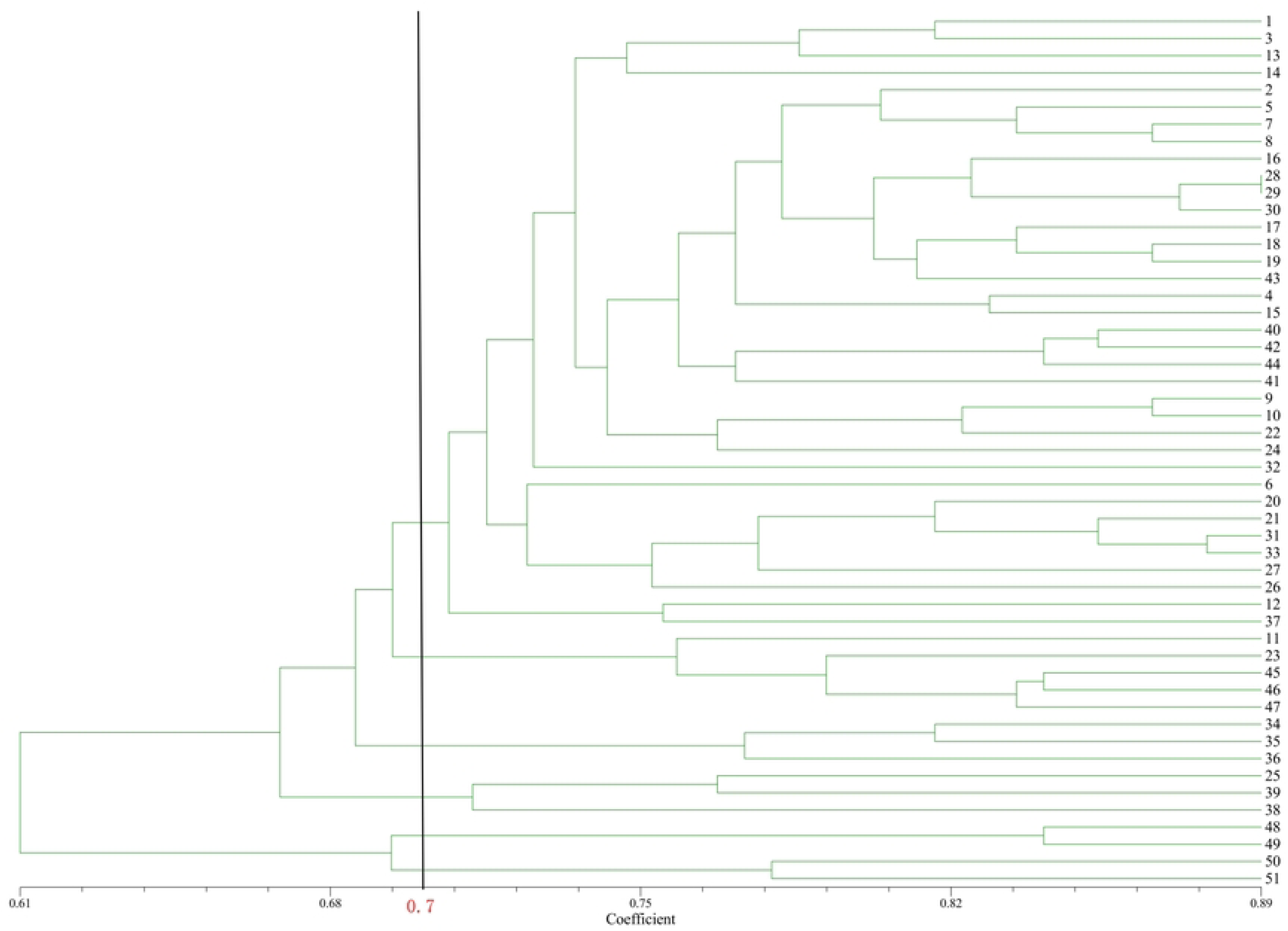
UPGMA clustering analysis diagram of 51 mulberry germplasm resources. **Note:** The UPGMA clustering analysis was performed using the NTSYS software to cluster 51 mulberry trees, dividing the mulberry population into 6 groups with a probability of approximately 0.7.

In order to better evaluate the correlation between the genetic diversity of mulberry germplasm and active substances, SPSS 27 software was used to perform cluster analysis on the 1-DNJ content of 51 germplasms (Fig 7). It can be divided into 2 major categories. The first category is composed of three small subcategories, including 45 mulberry germplasms. Among them, the 1-DNJ content of the first subcategory ranges from 0.4854 to 0.8070 mg/g, with an average of 0.6456 mg/g, and the overall DNJ content is low; the second subcategory has a variation of 0.8632 to 1.0624 mg/g, with an average of 0.9426 mg/g, the DNJ content is at a medium level; the third subcategory has a range of 1.1254 to 1.3122 mg/g, with an average of 1.1989 mg/g, and the content is medium to high; the second category includes 6 mulberry germplasm, with a range of 1.5628 to 2.5300 mg/g, and the average is 1.9724 mg/g, and the DNJ content is at the top level overall. Correlation analysis showed that the first largest category (36 accessions) of UPGMA clustering included 32 accessions in the first category of DNJ content clustering. Among them, 20 accessions in the second subcategory had a duplication rate of 90%. These mulberry accessions all had medium and low 1-DNJ content levels. Except for Sandaoheze, the third, fourth and sixth categories are all mulberry germplasms with low 1-DNJ content. The 1-DNJ content in the second category is generally at a moderate to upper level, with an average value of 0.9532 mg/g. Hanza No. 3 and Jingsang in the fifth category are both high 1-DNJ content germplasm, with an average value of 1.6988 mg/g.

**Fig 7.**
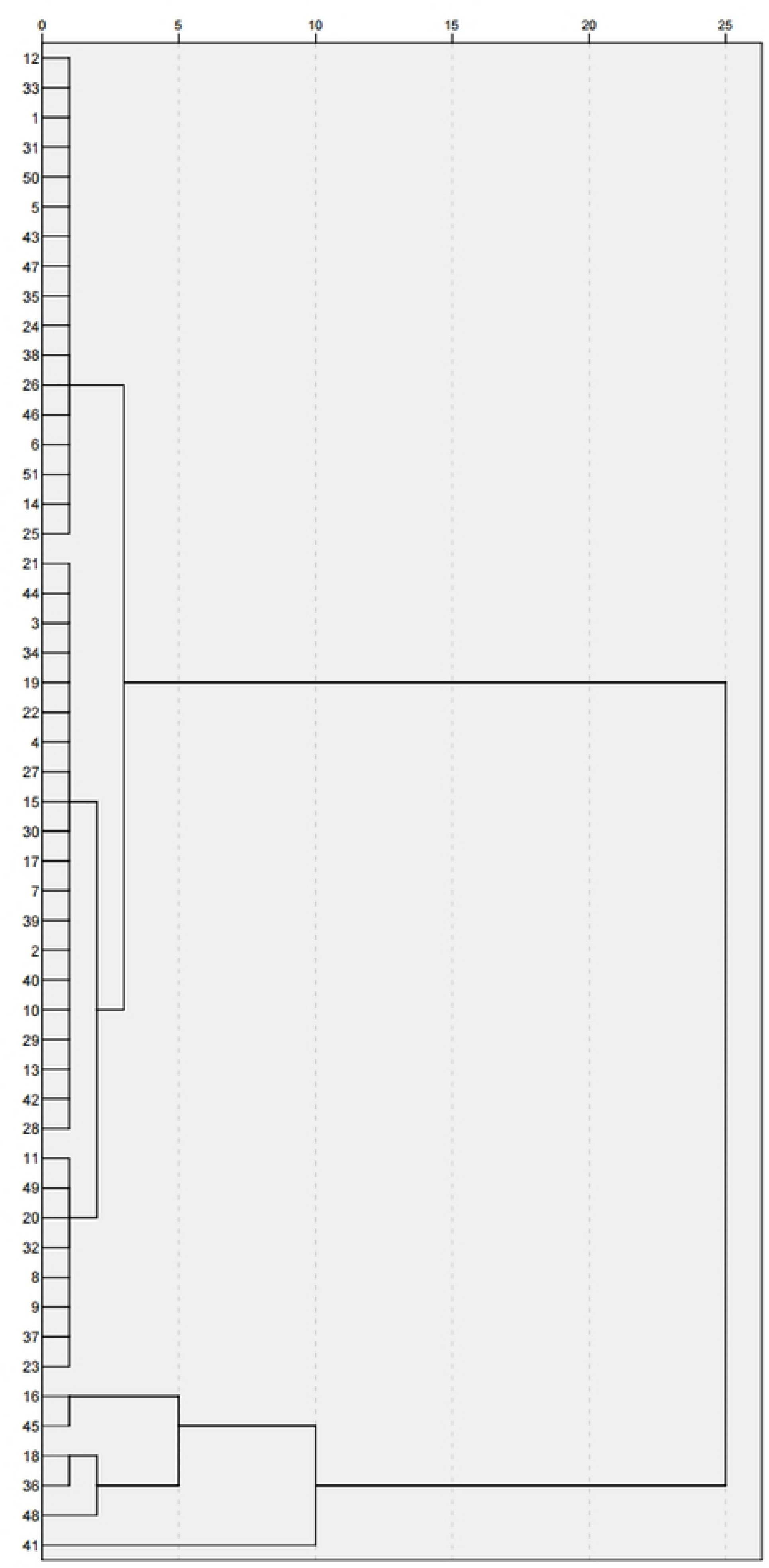
Clustering analysis of 51 mulberry germplasm resources based on leaf 1-DNJ content. **Note:** The phenotypic analysis results of 1-DNJ content in mulberry were performed using SPSS 27 software, and the mulberry population was divided into two major categories and four minor subcategories.

### Association analysis between SNP loci and 1-DNJ content

In order to evaluate the association between SNP polymorphic sites and 1-DNJ phenotypic variation, this study used 51 mulberry germplasms as objects, and conducted Mantel test on the 0/1 genotype matrix and 1-DNJ content of 81 polymorphic sites obtained through screening. The results showed that 6 of the 81 sites were significantly correlated with 1-DNJ content (p<0.05), and 2 of them reached an extremely significant level (p<0.01) (Fig 8). From the perspective of correlation strength, the signal of site Chr2-14 (1700 bp) is the strongest (r=0.2645, p<0.01), followed by Chr1-11 (400 bp) (r=0.2087, p<0.05) and Chr2-29 (260 bp) (r=0.1329, p<0.05); the correlation coefficients of the other three sites (Chr2-17 800 bp, Chr2-16 400 bp, Chr1-11 900 bp) ranged from 0.052 to 0.109, and the effect was relatively weak. The above six significant association sites can be used as candidate intervals for subsequent genetic analysis of 1-DNJ traits and functional gene mining.

**Fig 8.**
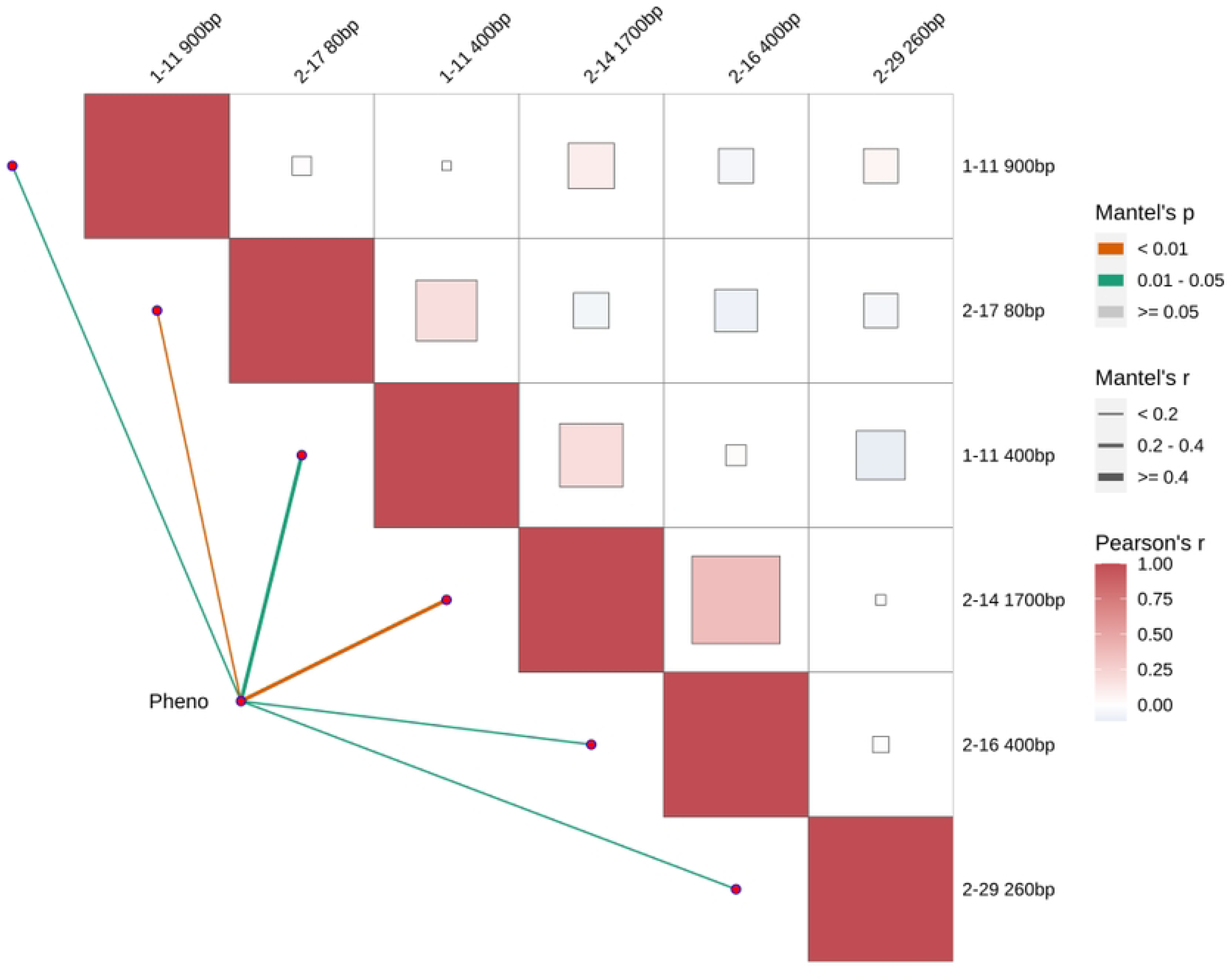
Mantel-test Heatmap of partial SNP markers and 1-DNJ content.

## Discussion

1-DNJ is the core hypoglycemic active component in mulberry and a key indicator for evaluating its medicinal value. The content of 1-DNJ in mulberry leaves is affected by germplasm resources, growth stages and cultivation conditions, exhibiting significant interspecific and intraspecific varietal differences[23]. The 1-DNJ content of the 51 mulberry germplasms assayed in this study ranged from 0.4854 to 2.5300 mg/g, with a coefficient of variation of 0.4241, which indicated a significant genetic differentiation in this trait. This result is consistent with the conclusions of previous studies that the DNJ content in mulberry germplasms is governed by genetic background, and further verifies the genetic stability and exploitable potential of 1-DNJ content as a heritable trait.The 1-DNJ content of Yu71-1, Hanza No.3 and Sandaoheze among these germplasms exceeded 2.0 mg/g, endowing them with extremely high potential for medicinal exploitation and thus serving as core parental materials for the breeding of mulberry varieties with high 1-DNJ content. Notably, the distribution of 1-DNJ content exhibited no obvious geographical clustering pattern; germplasms with high, moderate and low 1-DNJ content were distributed across all production regions. This phenomenon may be attributed to frequent germplasm exchange during the long-term artificial domestication of mulberry, and also implies that the genetic regulatory mechanism underlying 1-DNJ content may not be directly restricted by geographical environments.There are significant differences in 1-DNJ content in mulberry leaves among different germplasm resources. For example, Guisangyou 12 (0.6317 mg/g) and Guisangyou 62 (0.9621 mg/g) have higher 1-DNJ contents than those reported in a previous study[24]. In contrast, Longsang No.1 (0.5143 mg/g), Baiguo mulberry (0.9666 mg/g), and Autumn mulberry (0.9945 mg/g) have lower 1-DNJ contents than those in another study [25]. Additionally, Nongsang No.14 (1.3122 mg/g) and Yu 71-1 (2.5300 mg/g) have significantly higher 1-DNJ contents than those recorded in a recent study [26], while Hongguo No.3 (0.4854 mg/g) has a notably lower content than that in the same study. It should be clarified that although the mulberry germplasm tested in this study was all planted in the Shaanxi region, the 1-DNJ content still showed significant differences. These differences arise not only from the genetic background and physiological characteristics of the mulberry germplasm itself but also from the regulatory effects of cultivation conditions, such as climate factors including light, temperature, and humidity, as well as soil types, on the content of plant secondary metabolites[9]. This further explains why there are variations in 1-DNJ content among different germplasms within the same planting area and among the same germplasm in different studies, providing an important reference for subsequently regulating mulberry 1-DNJ content through a combination of genetic improvement and optimized cultivation practices.

In terms of SNP-PCR system optimization, this study determined the optimal 20 μL reaction system through a combination of orthogonal experiments and single-factor experiments. The system includes 2.2 µL of Buffer (containing Mg²^+^), 0.4 μL of 2.5 mM dNTPs, a total of 2.75 μL of forward and reverse primers (10 μmol·μL^-1^), 0.3 μL of Taq DNA polymerase (5 U·μL^-1^), 1.1 μL of DNA (50 ng·μL^-1^), and 13.65 μL of ddH₂O, which can achieve stable and efficient amplification.The advantage of this system lies in balancing reaction specificity and amplification efficiency. In validation, it exhibited clear bands and abundant polymorphism, with the percentage of polymorphic loci reaching 89.10%. This value was significantly higher than the polymorphism levels of traditional molecular markers such as SSR and ISSR reported in previous studies, including 71.79% (introduced germplasm) and 82.56% (wild germplasm)[27], an average polymorphism ratio of 76.67% [28], a polymorphism ratio of 78.57% [29], and a polymorphism ratio of 80.9% [30]. These results demonstrated that the SNP markers developed based on genome resequencing have higher resolution and are more suitable for the accurate analysis of genetic diversity in mulberry germplasm. Compared with previous studies, this system has optimized the primer dosage and template concentration, which reduces the risk of non-specific amplification and provides more reliable technical support for molecular marker-assisted breeding of mulberry.

Among the 23 highly polymorphic primers screened in this study, Chr2-35 (0.6681), Chr2-36 (0.6749), and Chr6-8 (0.6906) showed Nei’s diversity indices significantly higher than the average value (0.4667) and markedly higher than the genetic diversity levels of most primers. Based on the phenotypic data in this study, it can be preliminarily inferred the genomic region targeted by the above primers may be a candidate segment for subsequent mining of 1-DNJ related sites. Notably, the SNP sites corresponding to highly polymorphic primers such as Chr2-16 and Chr2-17 in this study are, according to functional predictions, located in specific gene coding regions, with the predicted functions of the genes at these two loci being endo-1,3;1,4-beta-D-glucanase and basic 7S globulin-like protein, respectively. Based on previous studies, SNPs in coding regions may lead to changes in the amino acid sequence of the corresponding proteins, thereby affecting their spatial structure and functional activity: endo-1,3; 1,4-beta-D-glucanase is a common glycoside hydrolase in plants, mainly involved in processes such as cell wall remodeling and defense responses[31]; basic 7S globulin-like proteins often serve as seed storage proteins, and some members can indirectly influence the synthesis and accumulation of secondary metabolites by regulating nutrient distribution in the plant [32]. It is inferred that changes in the functions of these two types of proteins may indirectly regulate physiological metabolic processes in mulberry, affecting the synthesis and accumulation of 1-DNJ. Furthermore, as a core medicinal active component in mulberry, the biosynthesis of 1-DNJ is a complex, multi-step enzyme-catalyzed process. Previous studies have shown that glycosyltransferases, lysine decarboxylases, and other enzymes are key in its biosynthesis [33]; Basic 7S globulin-like proteins are mostly used as seed storage proteins, and some members can also indirectly participate in the regulation of the synthesis and accumulation of secondary metabolites by regulating the distribution of nutrients in plants [34]. It is speculated that changes in the functions of these two types of proteins may affect the synthesis and accumulation of 1-DNJ by indirectly regulating the physiological metabolic process of mulberry trees. In addition, as the core medicinal active substance of mulberry, the biosynthesis of 1-DNJ is a complex process of multi-step enzymatic reactions. Previous studies have shown that glycosyltransferase and lysine decarboxylase are key enzymes in the synthesis process [31]. However, the genetic loci related to 1-DNJ content found in this study do not involve the above-mentioned key enzymes. This also suggests that there may be multiple pathways for the genetic regulation of 1-DNJ content in mulberry, which provides a new research direction for the subsequent in-depth analysis of the genetic regulation mechanism of 1-DNJ content in this study.

This study found that there is a certain correspondence between UPGMA clustering based on SNP markers and phenotypic clustering based on 1-DNJ content in the overall grouping pattern. UPGMA clustering divided 51 germplasms into 6 categories; DNJ content clustering divided them into 2 major categories. The duplication rate of the two types of clusters in the medium-low content germplasm is about 90%. Among them, UPGMA type I contains 32 accessions of the first type of 1-DNJ cluster. The average 1-DNJ content of type II germplasm is 0.9532 mg/g, and the average value of type V (Hanza No.3, Jingsang) is 1.6988 mg/g, indicating that there is a synergistic trend between molecular differentiation and phenotypic variation, but there is a deviation in the attribution of some germplasms in the two clusters, suggesting that the strength of the association between the whole-genome genetic variation reflected by SNP markers and the phenotypic variation of 1-DNJ is limited. The Mantel test further verified this trend. Among the 81 polymorphic sites, 6 sites were significantly correlated with 1-DNJ content, but the correlation coefficients did not exceed 0.3 (r=0.052∼0.265), which was a weak correlation. This result is consistent with the “limited synergy” phenomenon observed in the cluster comparison in this study, both indicating that the association between SNP markers and 1-DNJ content is real but weak. The reason for this weak correlation may be that the synthesis and accumulation of secondary metabolites in plants are usually jointly regulated by a complex network of multiple minor genes, rather than by a single major gene [35], and a single genetic locus has limited effectiveness in explaining phenotypic variation. In summary, this study identified excellent germplasm with high DNJ content, optimized the molecular marker system suitable for mulberry germplasm identification, systematically compared the clustering trends of molecular clustering and 1-DNJ phenotypic clustering, and preliminarily located the SNP sites linked to 1-DNJ content through Mantel test, which provides important scientific basis for the identification of mulberry medicinal germplasm, molecular-assisted breeding, and DNJ-related medicinal development.

## Conclusions

This study systematically determined the 1-deoxynojirimycin (1-DNJ) content in 51 mulberry germplasm resources, optimized the SNP-PCR reaction system, and analyzed their genetic diversity, aiming to provide a scientific basis for the breeding of medicinal mulberry varieties and the investigation of their genetic regulatory mechanisms. The results showed that the 1-DNJ content in the leaves of the tested mulberry germplasms ranged from 0.4854 to 2.5300 mg/g with a coefficient of variation of 0.4241, indicating significant genetic differentiation among the germplasms and thus laying a solid foundation for the screening of elite medicinal mulberry germplasms. An optimized SNP-PCR reaction system (20 μL total volume) for mulberry was successfully established, consisting of 2.2 μL of Mg²⁺ containing PCR buffer, 0.4 μL of 2.5 mM dNTP mixture, 2.75 μL of mixed forward and reverse primers (10 μmol·μL⁻¹), 0.3 μL of Taq DNA polymerase (5 U·μL⁻¹), 1.1 μL of genomic DNA template (50 ng·μL⁻¹), and 13.65 μL of sterile double-distilled water (ddH₂O). Validation experiments confirmed that this system exhibited excellent amplification stability and specificity, making it suitable for large-scale SNP genotyping of mulberry germplasms. Based on this optimized system, 23 pairs of high-quality polymorphic SNP primers were screened out from 72 primer pairs, which amplified a total of 91 loci across the tested germplasms, among which 81 were polymorphic. The percentage of polymorphic loci reached 89.10%, which was significantly higher than that detected by traditional molecular markers. Genetic diversity parameters (effective allele number 1.5362, Nei diversity index 0.4667, Shannon information index 0.3104) indicate that the test population has a moderate to upper level of genetic diversity. UPGMA clustering divided 51 germplasms into 6 major groups, which showed a certain degree of synergy with the phenotypic clustering (2 major categories and 4 subcategories) based on 1-DNJ content in the overall grouping pattern, but the correlation strength was limited. The Mantel test further verified this trend, in which 6 SNP sites were significantly but weakly correlated with 1-DNJ content (r<0.3, p<0.05), indicating that 1-DNJ, as a typical quantitative trait, may be slightly regulated by multiple genes. This study preliminarily identified excellent germplasm with high DNJ content and located the associated SNP sites. However, it only detected the 1-DNJ content at a single time point and failed to reveal its dynamic accumulation pattern. The functions of the candidate SNP sites need to be further verified by means of transcriptome or gene editing. In addition, the association analysis of 51 germplasms has limited effectiveness. Subsequently, the population size needs to be expanded and combined with genome-wide association analysis to more comprehensively analyze the genetic regulatory mechanism of 1-DNJ content.

## Author contributions

Conceptualized the study: Z.S. and J.L. Developed the methodology and software: Z.S. and Z.L. Performed validation and investigation: Z.S. and J.L. Conducted formal analysis: Z.S. and K.H. Provided resources: Z.S., Z.L. and J.G. Curated the data: Z.S. and J.L. Wrote the original draft: Z.S. Reviewed and edited the manuscript: J.L. Prepared the visualization: J.S. and F.W. Supervised and administered the project: J.L. Acquired funding: J.L. and J.S. All authors have read and agreed to the published version of the manuscript.

## Acknowledgements

We would like to express our sincere gratitude to all authors for their valuable contributions to the study design, data collection and analysis, as well as the preparation and revision of the manuscript.

## Conflicts of Interest

The authors declare no conflicts of interest.

## Data availability statement

The raw sequencing data supporting the findings of this study have been deposited in the NCBI Sequence Read Archive (SRA) under the BioProject accession number PRJNA1304463. These data are publicly available and can be accessed through the NCBI SRA database. The gel electrophoresis images and other supplementary materials are available in the PLOS ONE submission system as Supporting Information.

## Notes

### Competing Interest Statement

NO authors have competing interests Enter: The authors have declared that no competing interests exist.

